# Coronavirus, as a source of pandemic pathogens

**DOI:** 10.1101/2020.04.26.063032

**Authors:** T. Konishi

## Abstract

The coronavirus and the influenza virus have similarities and differences. In order to comprehensively compare them, their genome sequencing data were examined by principal component analysis. Variations in coronavirus were smaller than those in a subclass of the influenza virus. In addition, differences among coronaviruses in a variety of hosts were small. These characteristics may have facilitated the infection of different hosts. Although many of the coronaviruses were more conservative, those repeatedly found among humans showed annual changes. If SARS-CoV-2 changes its genome like the Influenza H type, it will repeatedly spread every few years. In addition, the coronavirus family has many other candidates for subsequent pandemics.

**One Sentence Summary:** The genome data of coronavirus were compared to influenza virus, to investigate its spreading mechanism and future status. Coronavirus would repeatedly spread every few years. In addition, the coronavirus family has many other candidates for subsequent pandemics.

## Introduction

COVID-19 (1, 2) is rapidly spreading worldwide. To investigate its spreading mechanism, genomes of the coronavirus were compared to those of the influenza virus (3) by principal component analysis (PCA) (4). These two RNA viruses differed in host specificity and speed of mutations. Here, I present the characteristics of both and the changes in coronavirus. The coronaviruses presented smaller variations and did not differ much from host to host. These characteristics may have eased the novel transfection of coronaviruses to humans; many of the bat strains seem to have the ability to infect humans, and the intermediate animals may not be necessarily required, against the present expectations. Although many of the coronaviruses were more conservative, their character may reflect a less selective pressure owing to their limited infectivity. Rather, those repeatedly found among humans showed annual changes. If SARS-CoV-2, which is highly infectious, changes its genome like the influenza H type, it will repeatedly spread every few years. In addition, the coronavirus family may have other candidates for subsequent outbreaks.

While the coronavirus genome is a positive-sense single-stranded 30 kb RNA, the influenza genome is divided in eight segments. Both directly replicate using their RNA-dependent RNA polymerases, which may cause many errors (5). This characteristic has introduced variations among coronaviruses, including the number and size of their open reading frames (ORFs). Some classes of coronaviruses, such as HCoV, cause upper respiratory tract infections in humans. The symptoms are mostly similar to those of the common cold, but they may also cause severe pneumonia (6). They have lower infectivity than the human influenza viruses. For example, a 2010-2015 study in China reported that 2.3% and 30% of patients were positive for coronavirus and influenza virus, respectively (6); a similar ratio was found in another large study (6). Some viruses may have a much higher infectivity and cause outbreaks: e.g., SARS-CoV (7, 8), MERS-CoV (7, 9, 10), and SARS-CoV-2 (SCoV2) (2, 11-13). The former two cause severe symptoms, while the latter varies from asymptomatic to critical.

The corona and influenza viruses have similarities and differences in infectivity, spread ability, and symptoms. These differences are based on their genomes, which are important for estimating how SCoV2 will act in humans.

## Materials and Methods

### Data and classification

Sequencing data were obtained from DDBJ database. Aligned data, obtained with DECIPHER(14) (presented in the Supplementary material), were further processed to observe the relationships among samples by using the direct PCA method (4), which can handle data with limited assumptions. The conceptual diagram of the PCA is as follows (all calculations were performed in R) (15). Updated versions of the scripts are presented in GitHub (https://github.com/TomokazuKonishi/direct-PCA-for-sequences). To escape the imbalance effect among samples, the decomposition was performed by removing clusters of similar samples, e.g. those caused by SARS, MERS, or SCoV2. Instead, only one sample was included from each cluster.

To prepare a comprehensive data set for SCoV2, 2,796 of full-length data were obtained from GISAID database and added to those used for Fig. 1. Some of those records were rather preliminary and contained several uncertain bases designated by “N”, which may be counted as indels. To cancel such artifacts, the corresponding regions were replaced with the average data in the PCA.

**Fig. 1.**
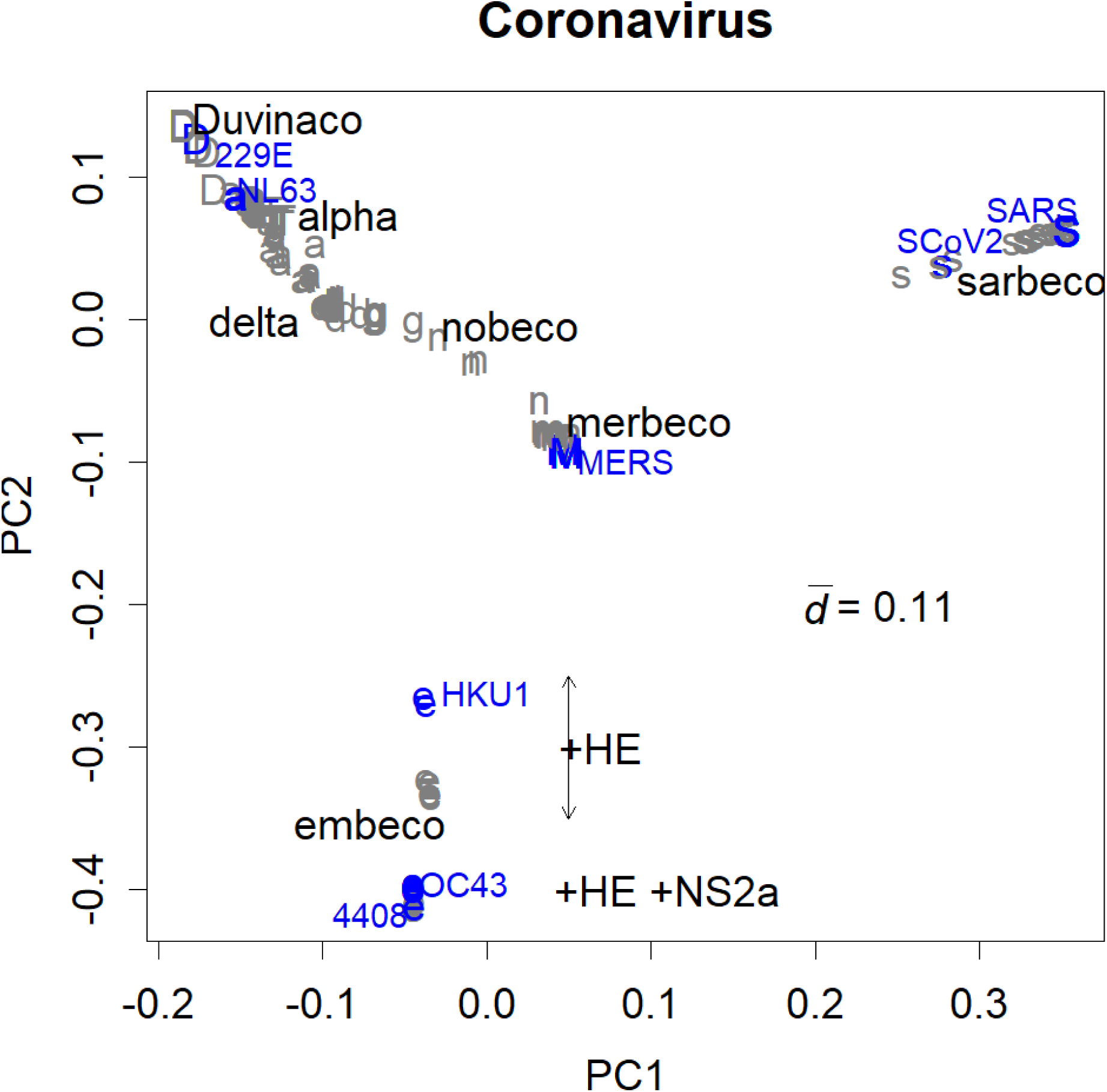
Classes of coronaviruses separated into PC1 and 2. Blue: human samples; subclasses of HCoV are indicated. a: Alphacoronavirus; d: Delta coronavirus; D: Duvinacovirus; e: Embecovirus; g: Gammacoronavirus; m: Merbecovirus; n: Nobecovirus; s: Sarbecovirus; T: TGEV. A major estimated class, Betacoronavirus, is not shown here, since the class has to cover distinctive classes (Embecovirus, Merbecovirus, Norbecovirus, and Sarbecovirus). It should be noted that many of the classification credits in the original record were different from this presentation.

### Diagram of the PCA

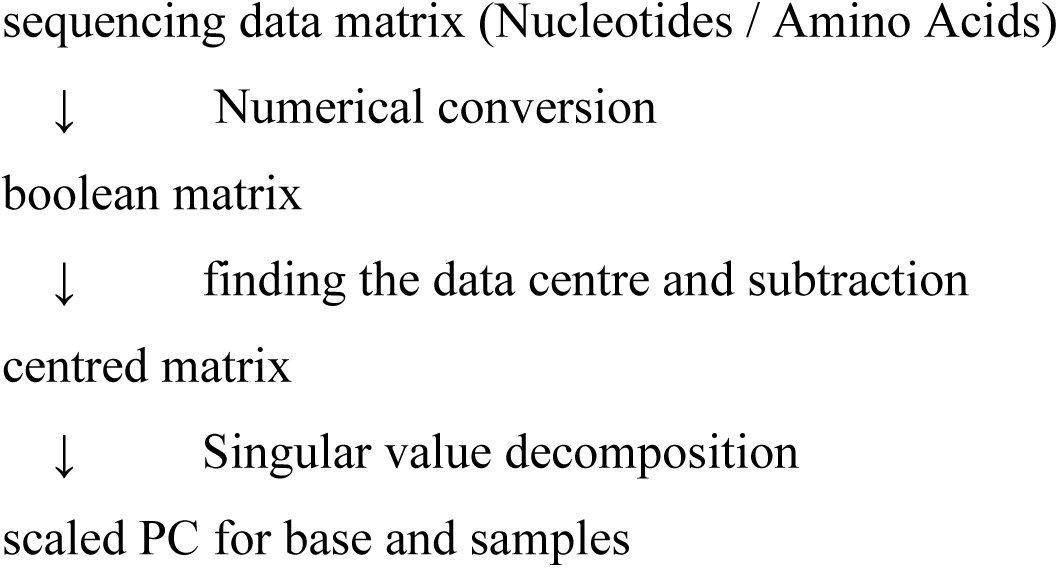

### Estimation of the magnitude of the sample variations

The scale among sample sequences was estimated by mean distances, scaled by the length of sequence *m*, of virus types. This was a type of standard deviation, 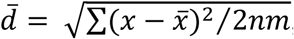, where *x*, 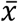, and *n* are the Boolean of each sample sequence, the mean sample, and the number of samples, respectively. Coefficient 2 was used to correct the double counts of the differences. The unit of length is the same as that of the PCA, which will extract the length toward particular directions.

### Estimation of mutation levels in the genome

The levels of PCs 1-5 for bases were estimated by the root sum square at each base position (Fig. 2 and S3). If alterations existed in several samples, and if they occur coincidently, they may contribute to a higher level of PC. To see the tendencies at the positions, two moving averages with a width of 200 amino acid residues were shown for substitutions (grey) and indels (blue).

**Fig. 2.**
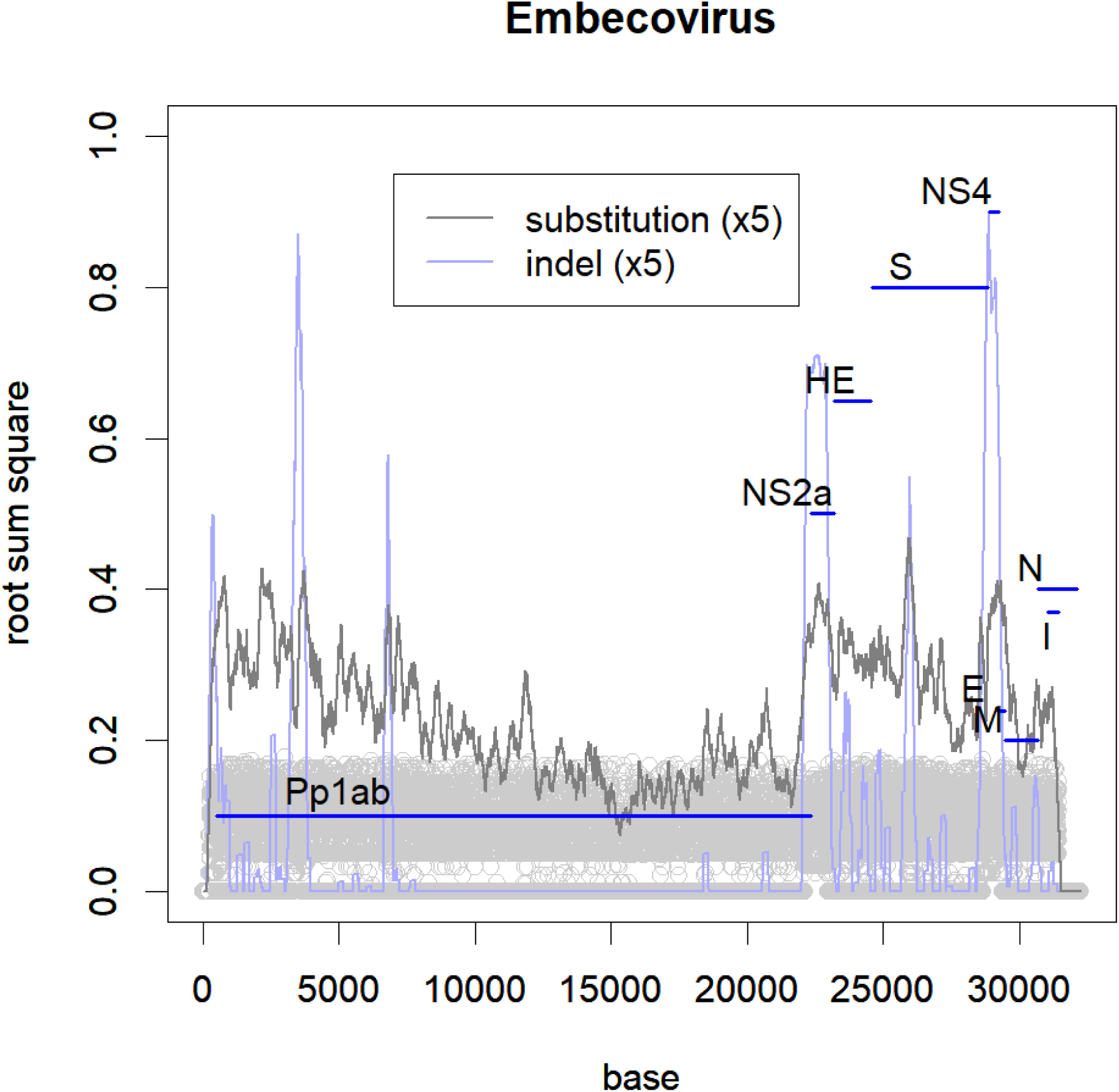
Frequency of mutations in each position of the genome. The level is estimated by the root sum square of PC1-PC5 (grey circles). The moving averages are represented in a 1:5 scale. The names of the ORFs are indicated. The indel at NS2a reflects that there are subclasses with the ORF and without the ORF.

## Results

The coronaviruses consist of distinct classes (Fig. 1). In the lower PC axes, the other classes were separated (Fig. S1a and b), and these were further divided into subclasses. For example, SARS-CoV and SCoV2 belong to different subclasses of Sarbecovirus (Fig. S1c and Table S1). The origin of the graph, (0, 0) coincides with the mean data. The accumulation of mutations will form a variety of viruses, which have different directions and distances from an original virus. If the mutations and samplings are random, the original virus would be near to the data mean. The variation magnitude, estimated by the mean distance 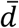, was 0.11. This is much smaller than those of single subclasses of influenza A virus (Fig. S2), such as H1 or H9. Incidentally, the value of 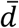 was not significantly altered by artificial reductions of sample numbers or sequence length (not shown).

Among the classes, Gammacoronavirus and Deltacoronavirus, which are close to the origin of the graph, were found mainly in bird samples (Fig. 1 and Table S1). These could be the origin of coronaviruses, as like the influenza viruses are thought to have originated from those of waterfowls. The mean of the studied samples was located in the bat virus Norbecovirus, which seems to be the origin of the viruses in mammals. Indeed, many classes apart from this class were found in bat samples. The most distant classes from the mean, TGEV and Embecovirus, were found in larger animals, but not in bats.

Human coronaviruses (HCoV) belong to Embecovirus, Alphacoronavirus, and Dubinacovirus (Fig. 1, blue). Two classes without such common cold viruses, Merbecovirus and Sarbecovirus, originated in recent outbreaks.

Similarly to other RNA viruses (5), many indels were observed, especially in some smaller ORFs (Fig. 2 and S3). These range from small regions without frameshifts to large ones that alter plural ORFs, e.g. Embecovirus is unique because it possesses an ORF of hemagglutinin. The class is further distinguished by having another ORF, NS2a, or not (Figs 1 and 2). Even within a small group of HCoV, OC43, an indel of 14 aa length existed in the spike protein. The classification was not significantly affected by either focusing on the indels or on the rest (Fig. S4). Therefore, indels were not given extra weight in this study; they were treated as a base or a residue. Note that some other small ORFs, such as the envelope and nucleocapsid, are conservative.

The values of PC were not significantly affected by the hosts (Fig. 1) e.g. differences between bird and swine viruses in Deltacoronaviruses were small (Table S1, PC18). This is contrarily to influenza viruses, which were separated among different hosts; the class for waterfowl is located near the centre, with three swine groups located around it, and two human groups were positioned in the most apart (Fig. S2a) (3). In coronaviruses, those apart from the Norbecovirus seem to infect larger animals, but this rule is not absolute (Table S1).

Each of the human-outbreak strains had similar ones in bats or camels, with minor differences (Fig. 1 and Table S1). In the SARS spike protein, no amino acid residue was unique to humans. This is partially because our knowledge about the viruses has increased after the efforts to screen likely viruses in wild animals (1, 16-18). Only 35 out of 2412 residues were different from SARS and similar bat viruses, and many of these were not conserved among the bat samples (Table S2). The situation was the same in SCoV2 which presented 34 unique amino acid residues (Table S2); however, this uniqueness could disappear after further research.

Influenza A H1N1 and HCoV yearly occurrence is very different, since only one H1N1 variety spreads worldwide yearly (3). Contrarily, several OC43 variants appear even within a single country (Fig. 3a, S5, and Table S3). H1 variants will never return in the subsequent seasons, whereas OC43 varieties appeared repeatedly for a decade. However, by concentrating solely on one variety, the annual alterations became obvious (Fig. 3b).

**Fig. 3.**
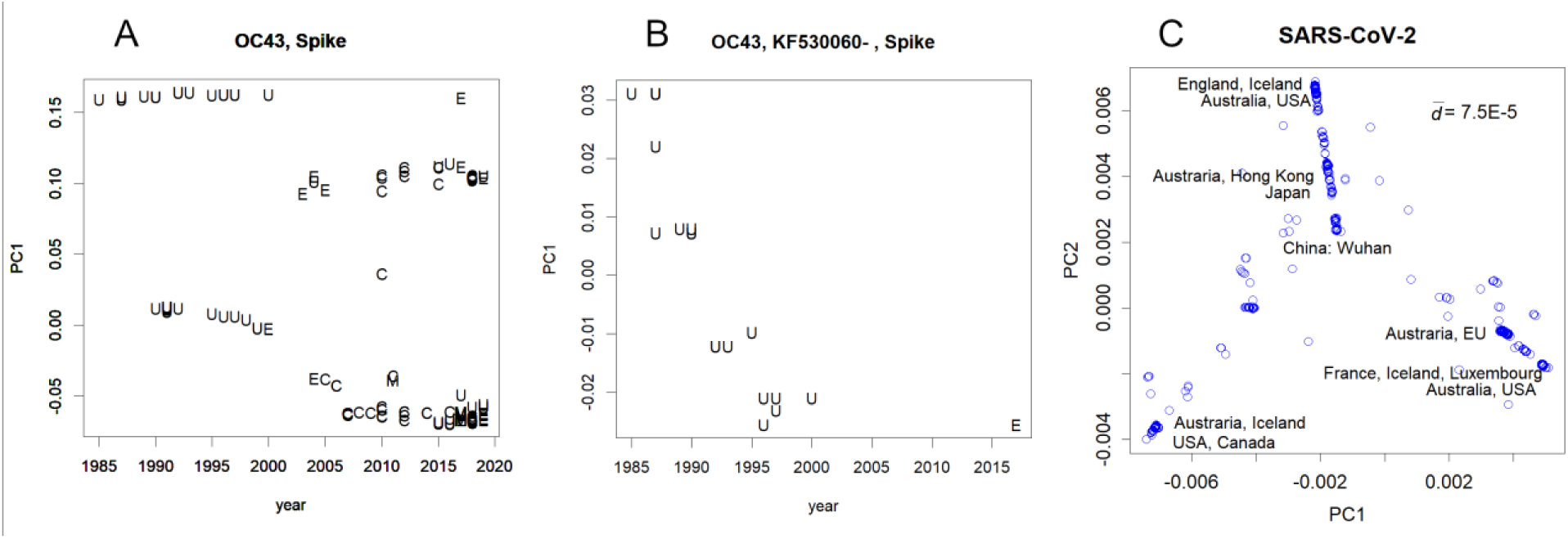
A. PC of the reported HCoV OC43 spike protein in each year. The letters represent the countries/regions: C- China, E- EU, M- Malaysia, U- USA. The upper-most series of variant were further selected to focus on them in the next panel. **B**, annual changes in the selected variant. Similar one-directional changes found in PC1 were always found in influenza H1N1 human cases ^3^. **C**. Classes of SCoV2 separated in PC1 and 2. Data from 2,836 samples are shown. Examples of countries are indicated. The full set of records is presented in Table S4.

A comprehensive set of SCoV2 samples was separated into three directions, forming some classes (Fig. 3c, S6a). Such classes could be made if a few persons with mutated virus migrated to another place and the virus started to infect people. Plural classes were found in some countries, suggesting multiple influx routes. The class closest to the data mean was that of China. Samples reported from the countries far from China tend to show larger magnitudes of PCs, and vice versa (Table S4).

Shift-type alterations were observed in coronaviruses, even though the genome is not separated. This is contrastive to the cases of influenza, which can replace RNA molecules between viruses. By focusing on the spike protein, coronaviruses were separated (Fig. 4a) similarly to the classification obtained by the whole genome (Fig. 1). However, in the classification obtained by 1ab polyprotein, the positions of Deltacoronavirus and Embecobirus were exchanged (Fig. 4b). Additionally, the position of the nucleocapsid protein in SARS-CoV moved from OC43, losing the Emvecovirus unity (Fig. 4c). These drastic changes are difficult to explain without shifts.

**Fig. 4.**
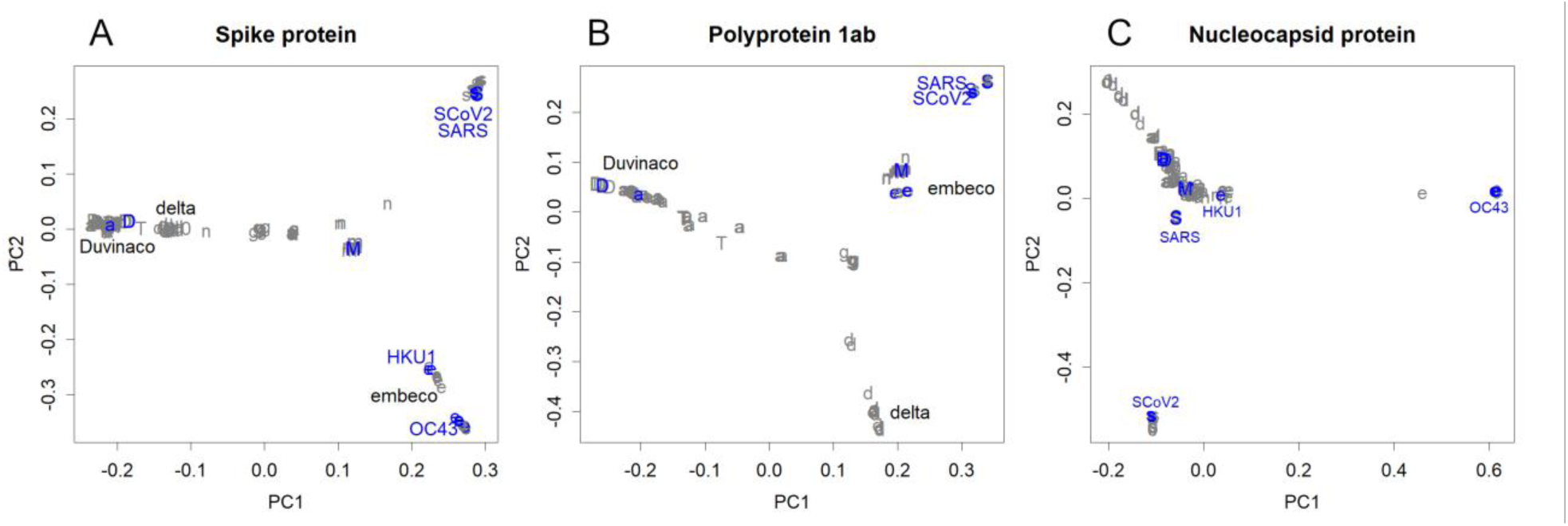
A Classification obtained from the amino acid sequences of the spike protein The relationships between the classes were similar to those estimated from the entire nucleotide sequences (Fig. 1). **B**. Classes found in polyprotein 1ab. The positions of the Deltacoronavirus and the Sarbecovirus were replaced. **C**. Classes found in the nucleocapsid protein. SARS-CoV moved out of the Sarbecovirus class.

## Discussion

Coronavirus and influenza classes were fairly different in the following two aspects: although the groups were clearly separated, the differences did not match those of the hosts in coronavirus; and the divergence magnitude among coronaviruses was much lower than that of a subclass of the influenza A virus. These characteristics corroborate the assessment that “coronaviruses can apparently breach cell type, tissue, and host species barriers with relative ease” (19). Coronavirus spike protein infection would be more tolerant to differences in hosts than the influenza virus’ hemagglutinin protein, which may have eased the selective pressure to separate the groups according to the various hosts.

Shifts, which are not possible by replacing virus segments, may have occurred in coronavirus. Since it does not copy the genome into double-stranded DNA, RNA splicing in nucleus (20) or RNA interference would be used instead of the ordinal homologous recombination by the RecA family (21). Those may also be the cause of frequent indels.

Viruses change their genome according to various selective pressures (5), such as to: a) Maintain the functions: any changes can cause malfunctions, hence they have to conserve the genomes, which is required for any state of the virus. b) Escape the herd immunity: Influenza A viruses, which are highly infectious, escape herd immunity by continuously changing. All ORFs change at the same speed (3), as any part of the virus that could be represented by the major histocompatibility complex will become a target of the immune system. Contrarily, HCoV is not very infectious; hence, some people remain unimmunized, easing this pressure. Also, the resulting smaller number of replications per year would slow the speed rate of changes. This might be the case for MERS-CoV among camels also (9, 22, 23). c) Exploit new hosts: this may require a change in the docking system. Adaptation to the genetic system of a new host may alter codon usage and several amino acids. SARS-CoV and SCoV2 might be under this type of pressure during infection. d) Increase asymptotic patients: patients with mild or no symptoms are required. In humans, once all infected individuals are identified, the virus is contained, especially if the symptoms are critical.

The conditions required to cause a pandemic are obvious in cases of influenza viruses. First, it has to be highly infective such as type H1N1. Second, it should be free from herd immunity. For example, the Pdm09 belongs to one of the two subclasses that did not cause outbreaks among humans (3). The SARS- and MERS-CoV fulfil these conditions, but they failed to escape the selective pressure mentioned above (d). SCoV2 satisfies all these conditions, thus spreading worldwide as Pdm09 did.

The dominated R type H1 of influenza A changed annually during its outbreak periods (3); by changing its most variable residues during three decades, and Pdm09 also changed annually. Coronaviruses have shown few annual changes (Fig. 3b and S6a), which might be due to their limited infectibility (HCoV) or to the lack of infected people (MERS-CoV). SCoV2 will face the selective pressure (b) as influenza A did. If it escapes this selective pressure, it will remain among humans and spread every few years. Actually, the change in SCoV2 has begun; they have formed several classes within the short emergence time (Fig. 3c, 6b). The magnitudes of the PC may show the migration pathways of the classes. They might mutate within China and transferred to other countries, and mutate further (Fig. S6b). The changes could be acclimation to humans (c); however, they may also relate to the herd immunity (b) and/or lower lethality (d). Fortunately, the lifespan of the classes of coronaviruses should be shorter than that of the influenza virus. The ORF lengths for the influenza virus are within a certain range, but some of the ORFs of coronavirus are quite short, e.g. the envelope protein, which is located in the conserved outermost region of the virus (24) (Fig. 2 and S3). It seems that this protein is too short to form a variable structure. Therefore, this will be a good target for herd immunity and these conservative ORFs might be suitable to produce vaccines. In contrast, the Spike protein tends to change (Fig. S3a, S3b), and may cause antibody-dependent enhancement (25).

Many bat coronaviruses seemed to be able to infect humans. The bat and human viruses are similar (Fig. 1) and there are more variations in bats, of which only a part we have observed. Due to replication errors and RNA editing, a bat may possess several virus variants (5). Although host-virus specificities are shown in laboratory experiments (19), as both humans and other animals have individual variations, the barrier would be more fragile in reality. Once an infection occurs, the virus will adapt through mutations, as for the SARS virus (18). Long-term accumulation of mutations in intermediate hosts, such as swine for influenza viruses, is not essential. These viruses would have a limited infectious character and cause mild symptoms to bats or other hosts; however, they could show an excessive adaptation to humans as SARS- and MERS-CoV did. As humans do not have herd immunity to many classes of coronaviruses (they do not contain HCoV), these might produce a new pandemic.

If intermediates are required, its main contribution could be the amplification of the inoculum size and contact frequency. Bats and human habitats are different, and a bat may not egest enough viruses to infect different hosts. The size of the donor animal is important, e.g. healthcare workers with secondary MERS infections tend to have milder symptoms and a better prognosis (23). This could be caused by differences in inoculum sizes; as camels produce plenty of nasal secretions full of viruses (22, 26), the inoculum size would be larger than that from humans. To prevent the production of intermediate animals, live animals should not be kept in the same place. Additionally, identifying the first patients is essential to prevent a human outbreak. Thus, people who frequently contact with living wildlife should not stay in a cosmopolitan city. The conventional classification system of coronavirus did not coincide with the relationships of the sequence data; e.g. the categories of Alpha- and Beta-coronavirus were too wide. Additionally, many of the credits for classification of original sequencing records were misjudged as well as those of the influenza virus. Using an objective method is preferable to determine the attributions (4).

## Abbreviations

HCoV: human coronavirus;
MERS: Middle East respiratory syndrome;
ORF: open reading frame;
PCA: principal component analysis;
SARS: severe acute respiratory syndrome;
SCoV2: SARS-CoV-2;
TGEV: transmissible gastroenteritis virus of swine

## Acknowledgments

The author would like to thank Editage (www.editage.com) for English language editing.

## Competing interests

The authors declare no competing interests.

## Data and materials availability

All data is available in the External Databases.

## Supplementary Materials

Figures S1-S6

Tables S1-S4

External Databases S1-S3

(an HTML version, which is easier to see, of the supplementary information is available in https://www.biorxiv.org/content/10.1101/2020.04.26.063032v2)

## Notes

### Competing Interest Statement

The authors have declared no competing interest.

### Summary of Updates

Aligned sequences in GISAID database have been removed. Please download them directory from the database.

