## Supplementary Figs and Tables for "Coronavirus, as a source of pandemic pathogens": index.htm

Supplement

### Supplementary Data

for **Coronavirus, as a supplier of pandemic viruses**  
Tomokazu Konishi  
  
Click to enlarge

- Fig. S1
- Fig. S2
- Fig. S3
- Fig. S4
- Fig. S5
- Fig. S6
- Table S1
- Table S2
- Table S3
- Table S4
- The corona virus sequences
- HCoV OC43 sequences
- SARS-CoV-2 sequences

#### Fig. S1

| a | b | c |
| --- | --- | --- |

  

Fig. S1. **a** and **b**. Classes of coronaviruses at the presented PCs. Blue: human samples. Labels are the same as Fig. 1. **c**. Classes found in the Sarbecovirus. SARS-CoV and SCoV2 belonged to different groups.

  

#### Fig. S2

| a | b | c |
| --- | --- | --- |
| d | e | f |

  

Fig. S2. Subclasses found in the indicated classes of the Influenza A virus. **a**. H1 hemagglutinin, **b**. H4 PB1, **c**. H5 hemagglutinin, **d**. H7 PB2, **e**. H9 hemagglutinin, **f**. H9 PB1. Values of the mean distance were indicated. The subclass may coincide with the hosts (**d**) but in many cases, a class was occupied by the limited host.

  

#### Fig. S3

| a | b | c | d |
| --- | --- | --- | --- |

  

Fig. S3. Levels of PC1-PC5 at each position in the nucleotide sequences. **a**. Sarbecovirus, **b**. HCoV OC43, **c**. MERS-CoV. **d**. SCoV2. Names of the ORFs are indicated as in Fig. 2.

  

#### Fig. S4

| a | b | Fig.1 |
| --- | --- | --- |

  

Fig. S4. Separation of classes in PC1 and 2, estimated by using indels (**a**) and substitutions (**b**).

  

#### Fig. S5

| a | b | c |
| --- | --- | --- |

  

Fig. S5. Subclasses of OC43 separated by the indicated PCs found in the whole genome. Conservative characteristics of the virus and repeated appearance were obvious.

#### Fig. S6

| a | b |
| --- | --- |

  

Fig. S6. **a** Annual changes in the MERS-CoV genome. **b** Comprehensive data for SARS-CoV-2. Samples found in some European countries showed higher magnitudes of PCs, indicating accumulation of mutations (also see Table S4).

  

#### Table S1

PC for samples of whole genome sequences of coronaviruses. Figure 1 was made from a part of this table.

#### Table S2

Differences in spike proteins a. SARS and bat SARS-like viruses. b. SCoV2 and other coronaviruses. Aligned protein sequences were compared. The IDs for the samples are listed.

#### Table S3

PC for samples of HCoV OC43 coronaviruses.

#### Table S4

PCs of SARS-CoV-2.

  
