## Supplementary figures and images for "Coronavirus, as a source of pandemic pathogens"

### Fig1.png

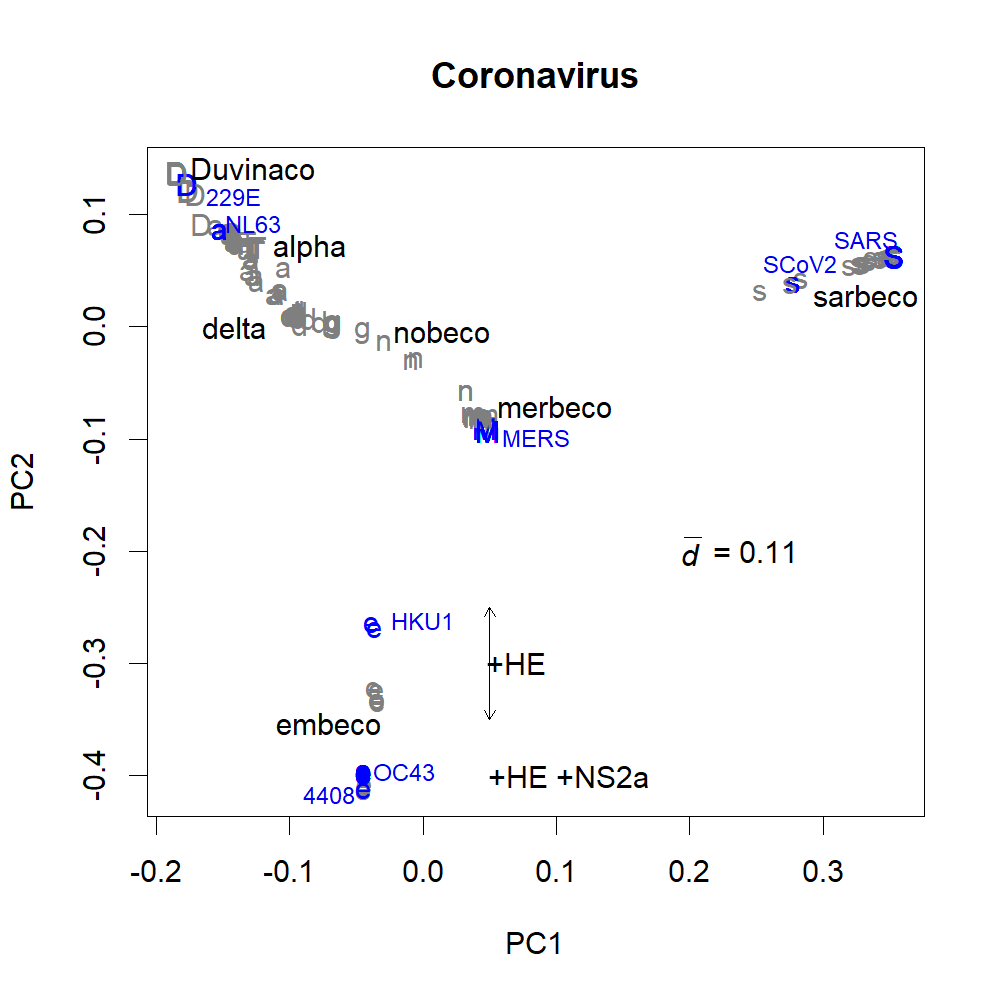

### S1a.png

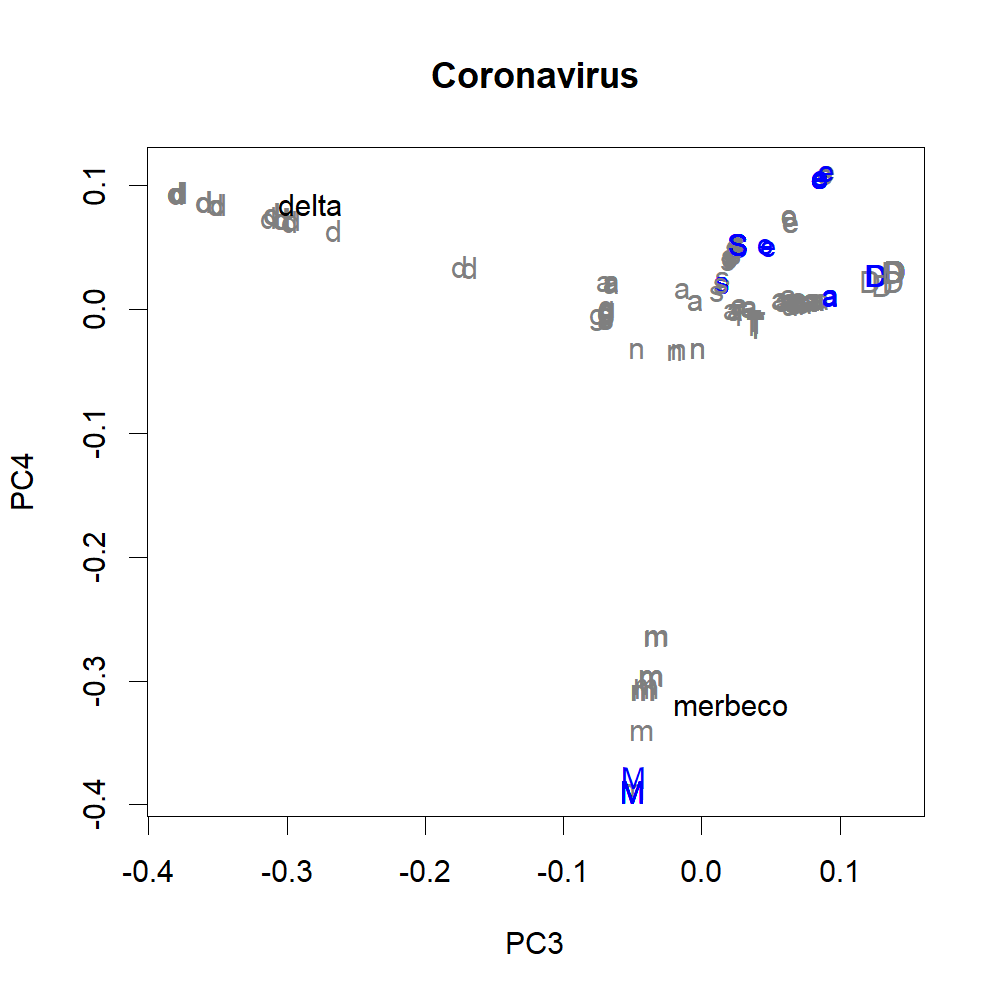

### S1b.png

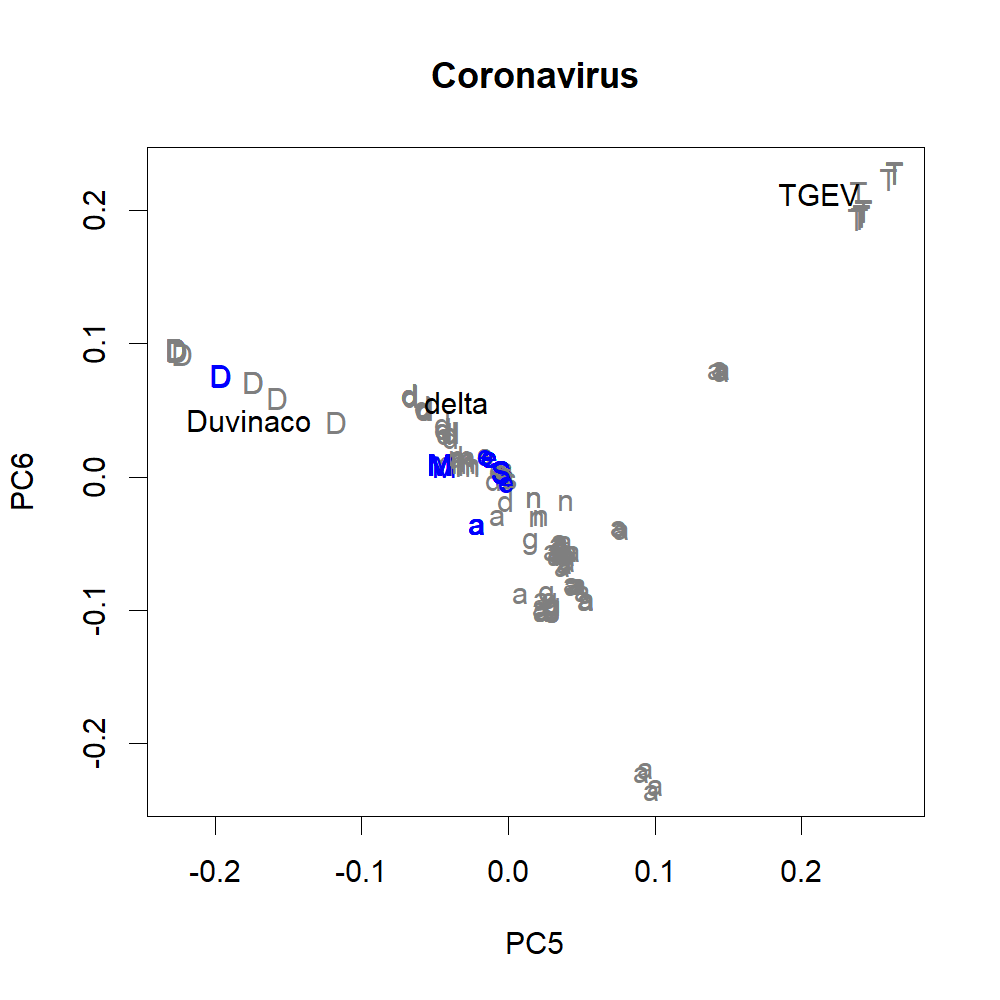

### S1c.png

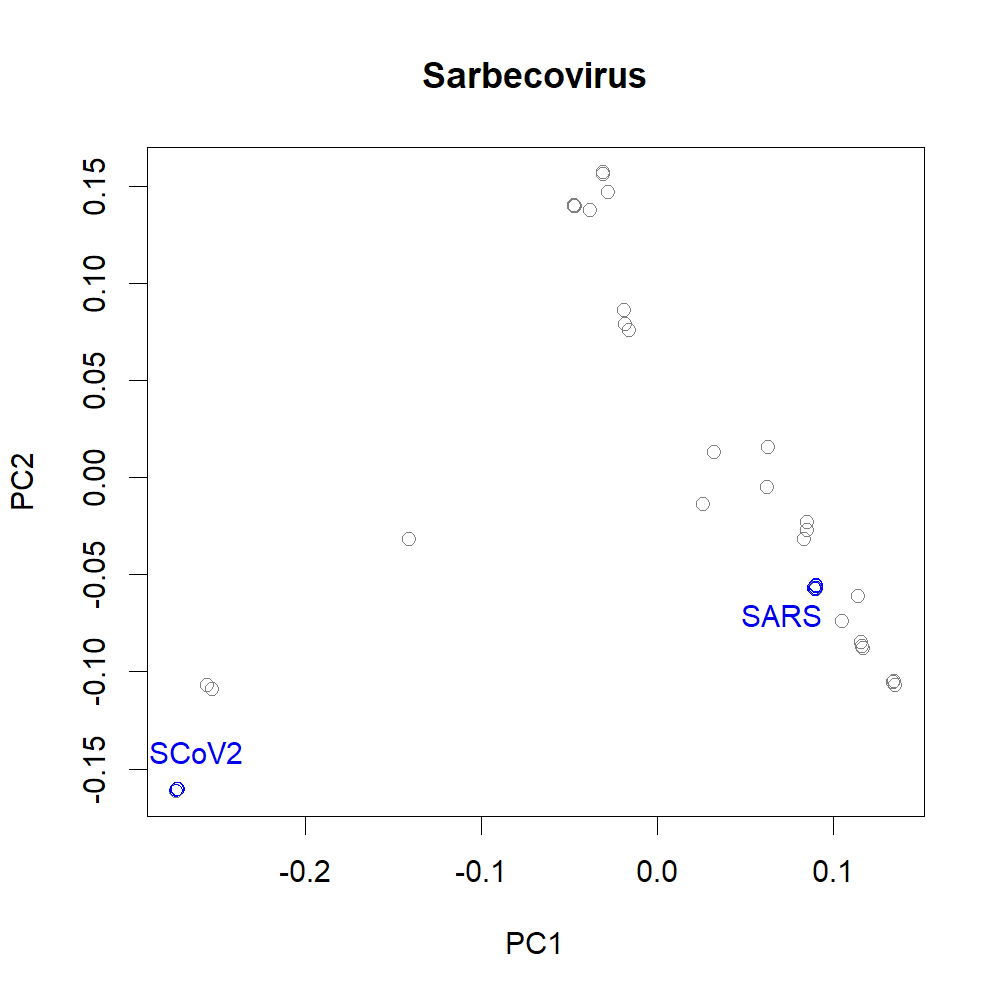

### S2a.png

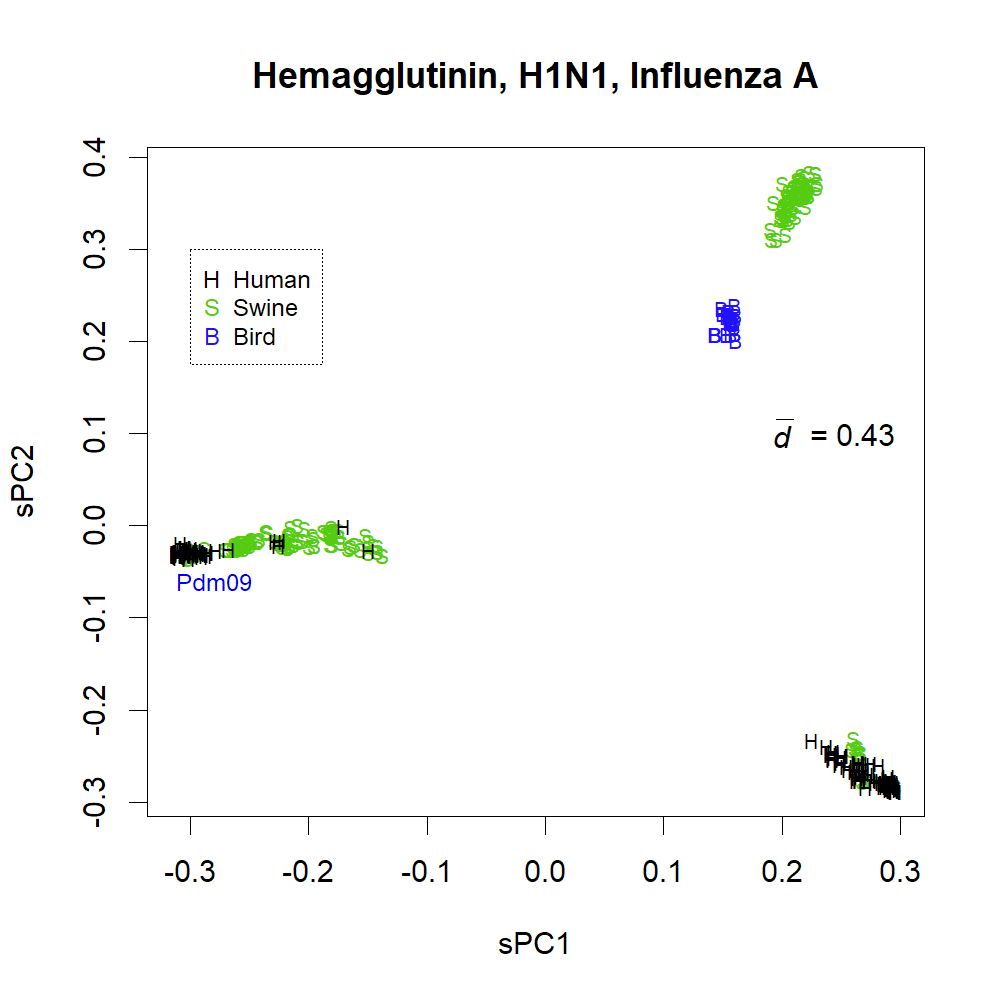

### S2b.png

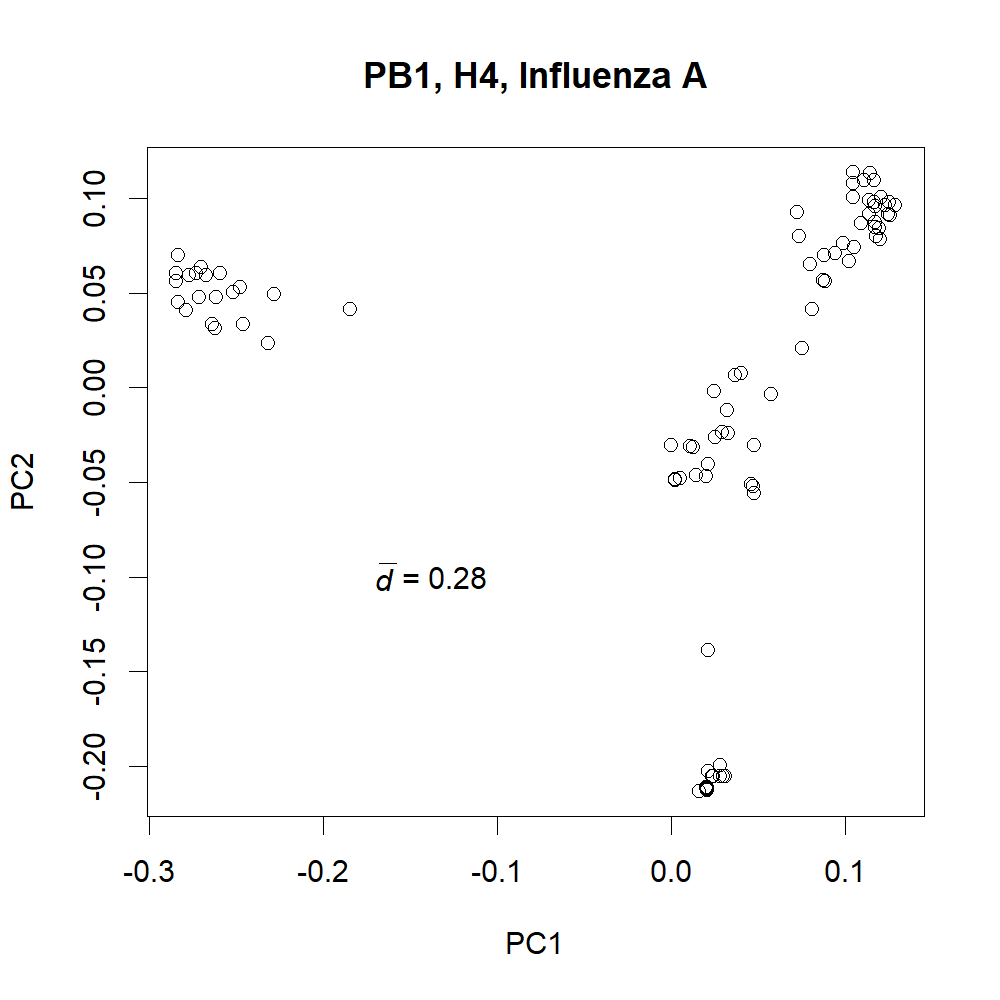

### S2c.png

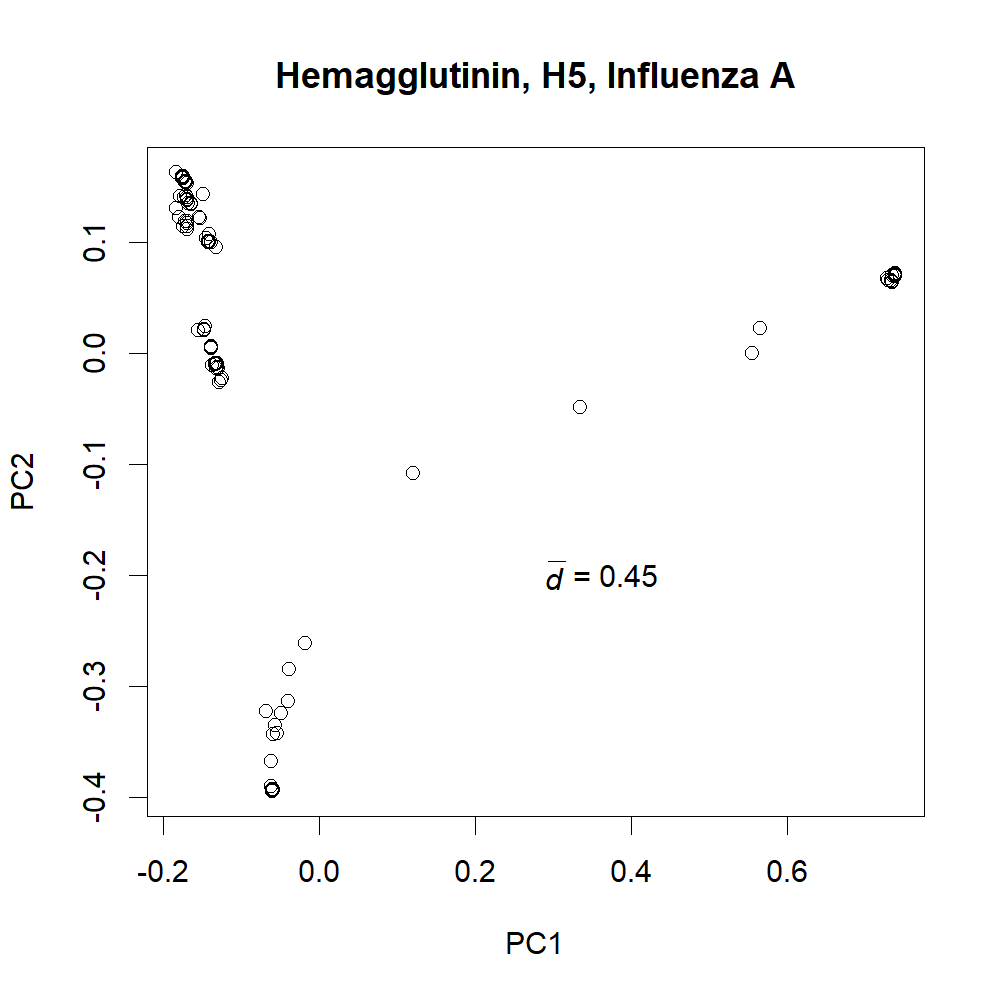

### S2d.png

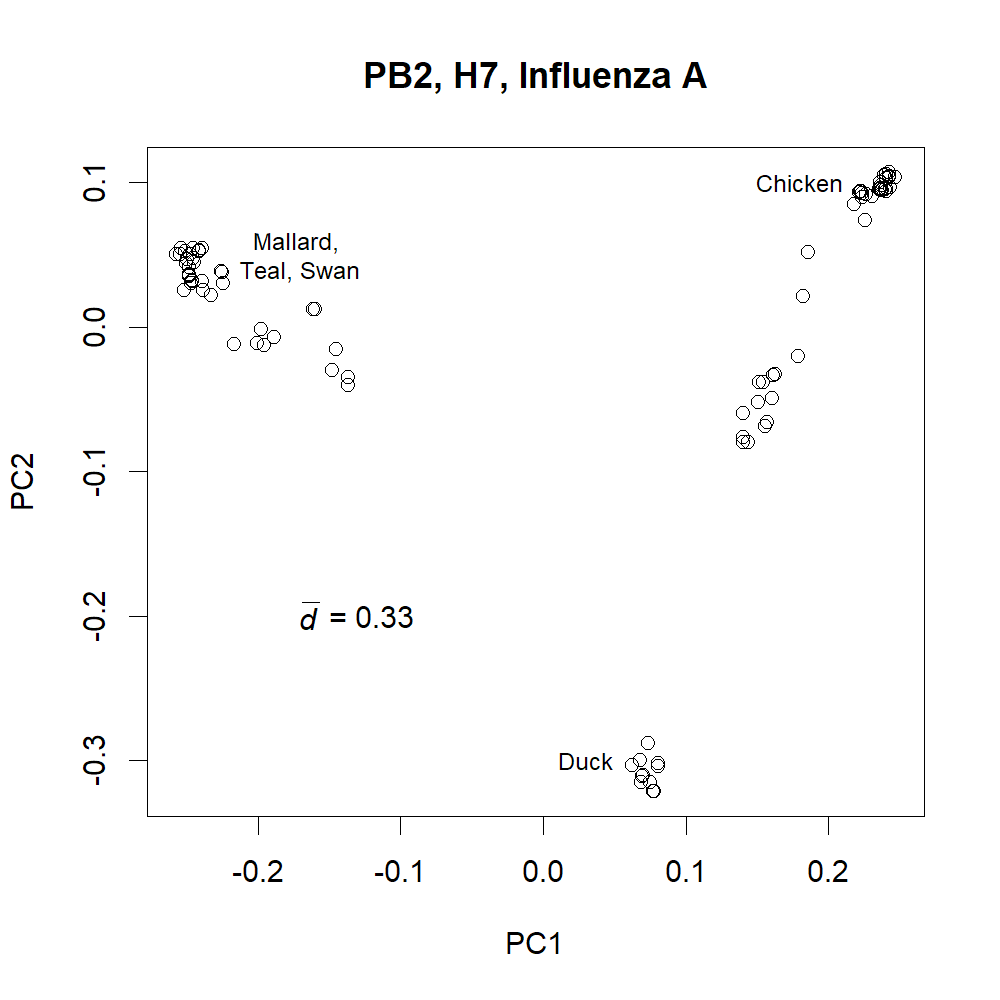

### S2e.png

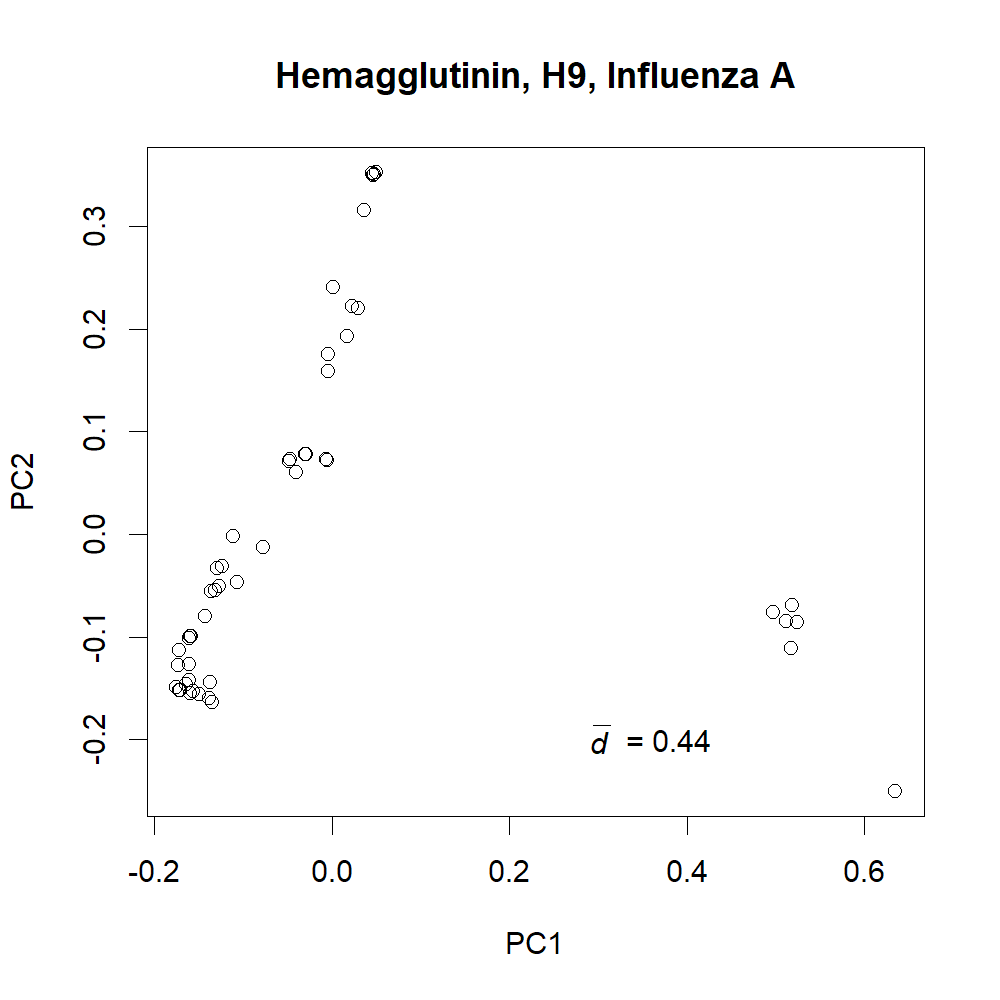

### S2f.png

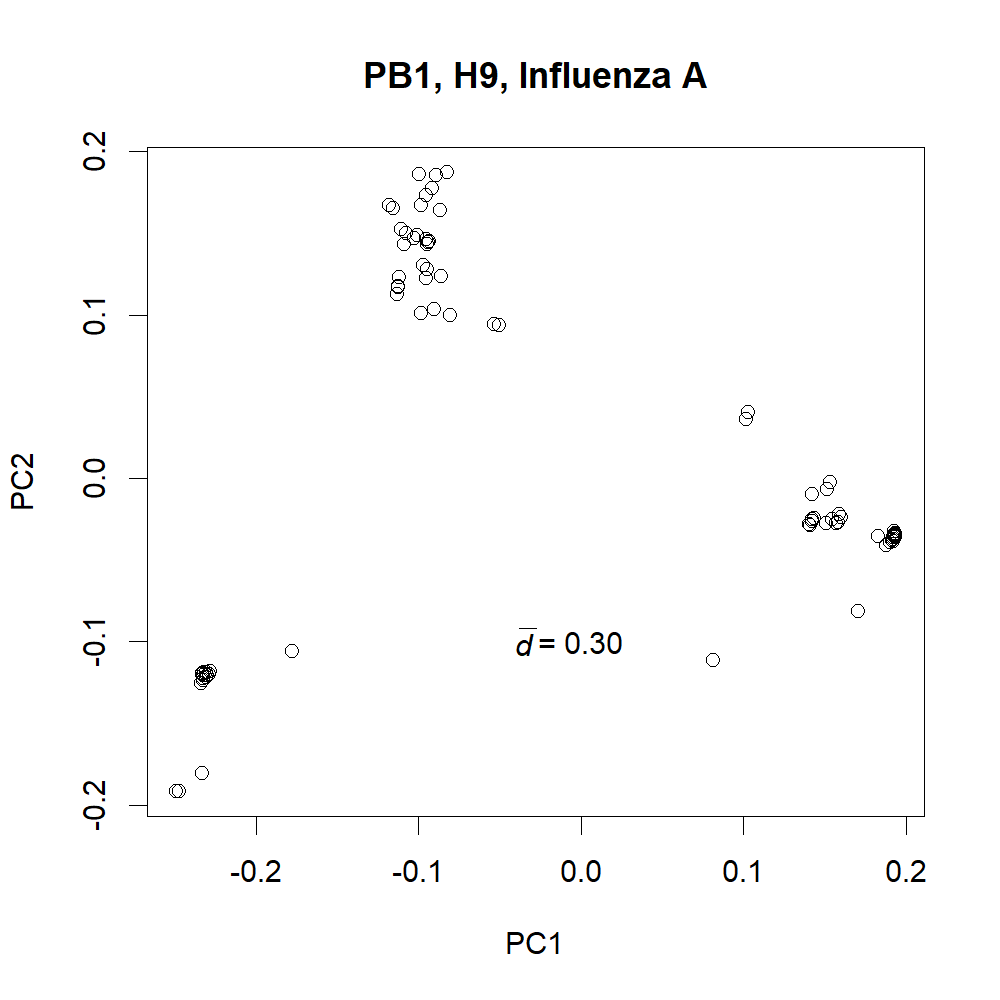

### S3a.png

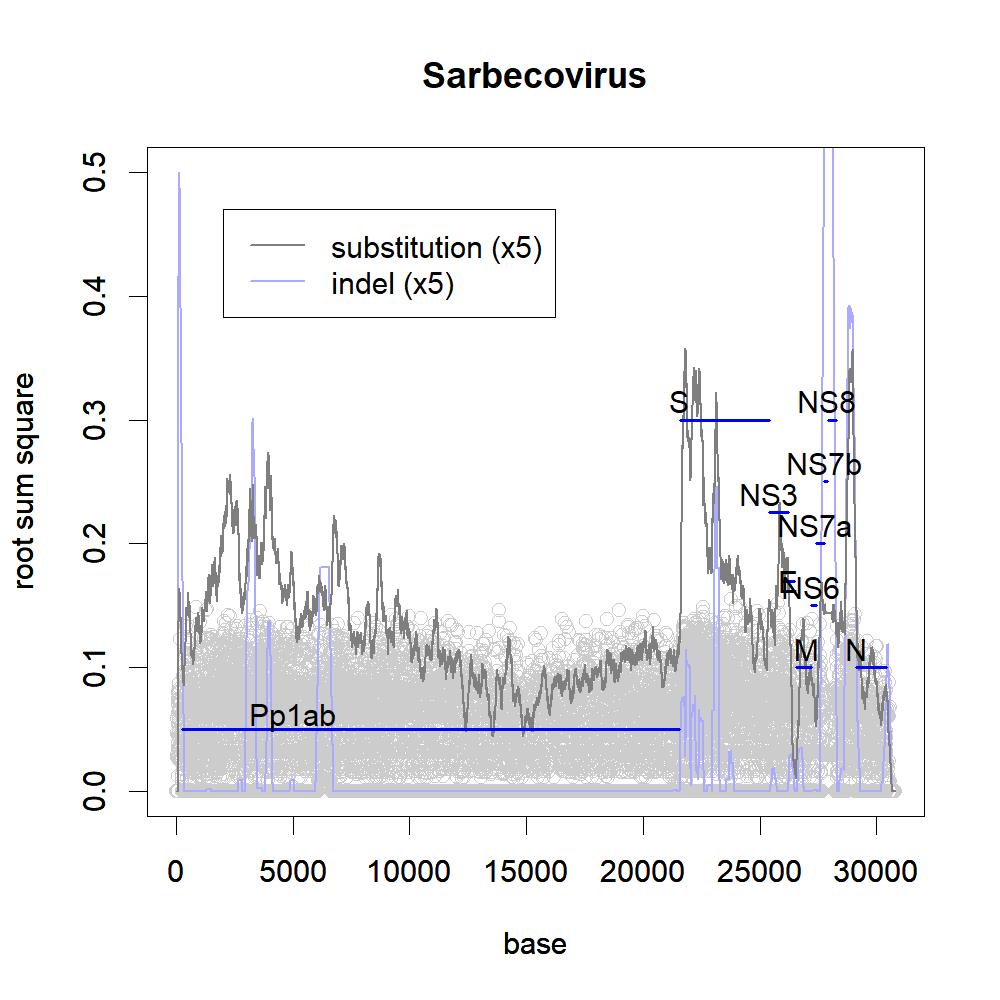

### S3b.png

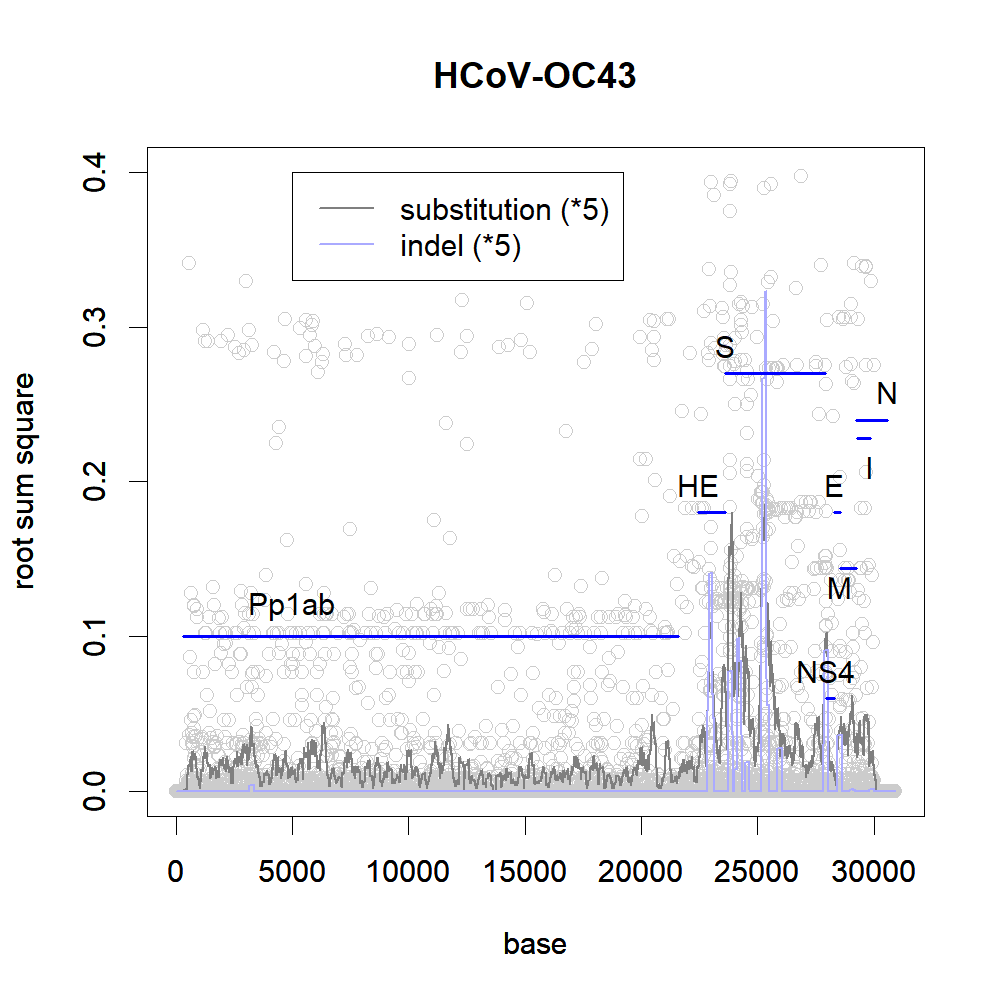

### S3c.png

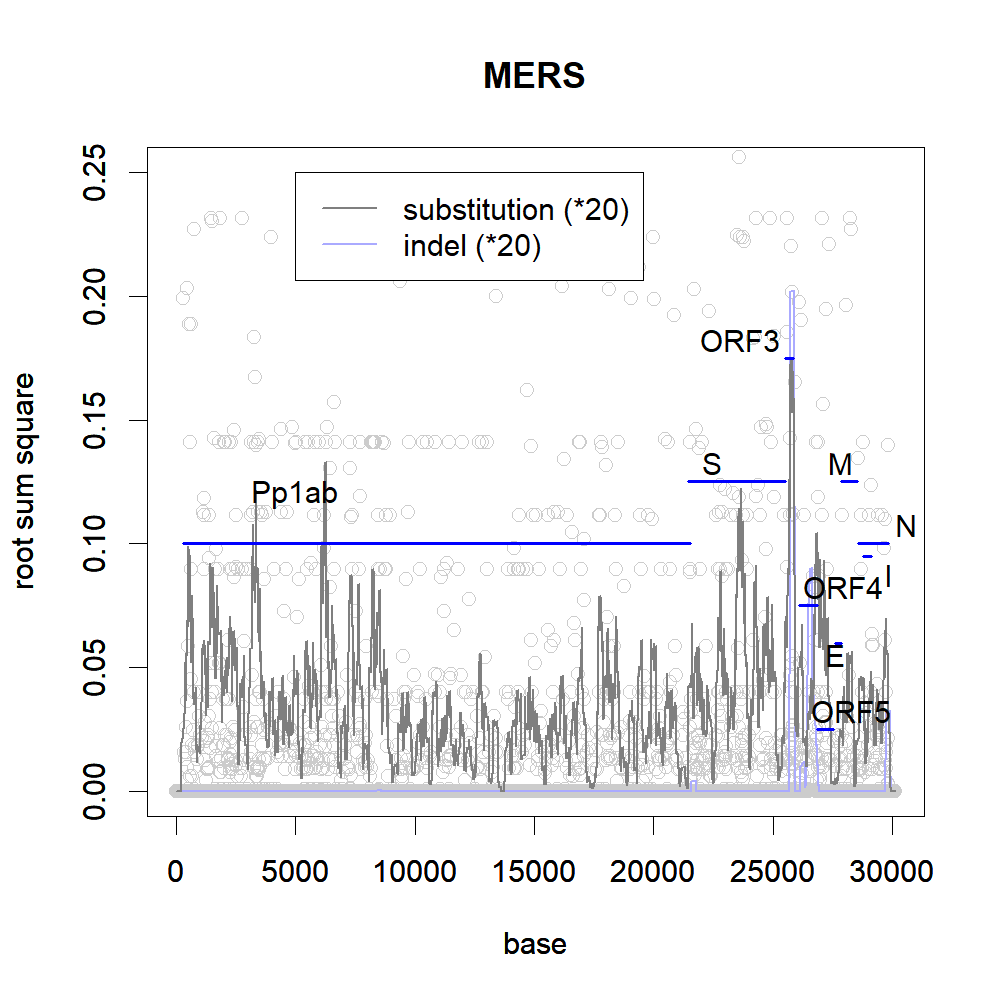

### S3d.png

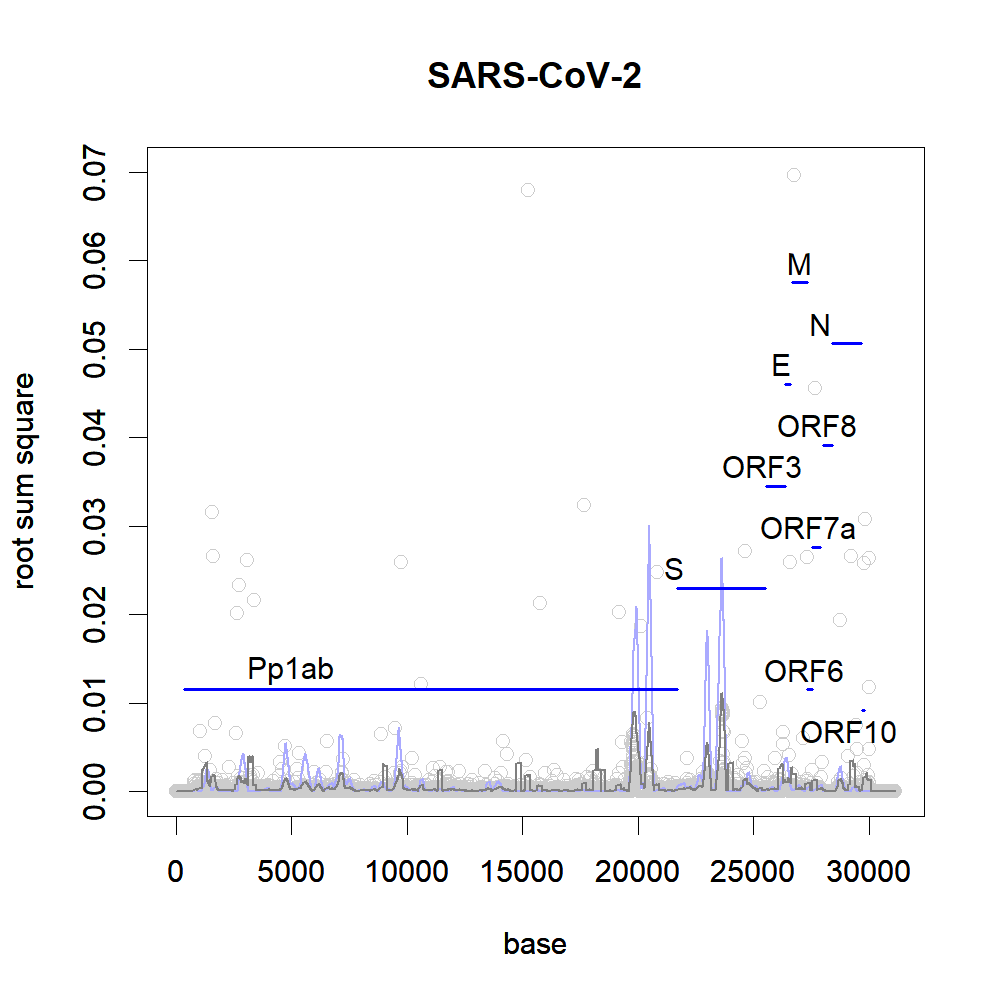

### S4a.png

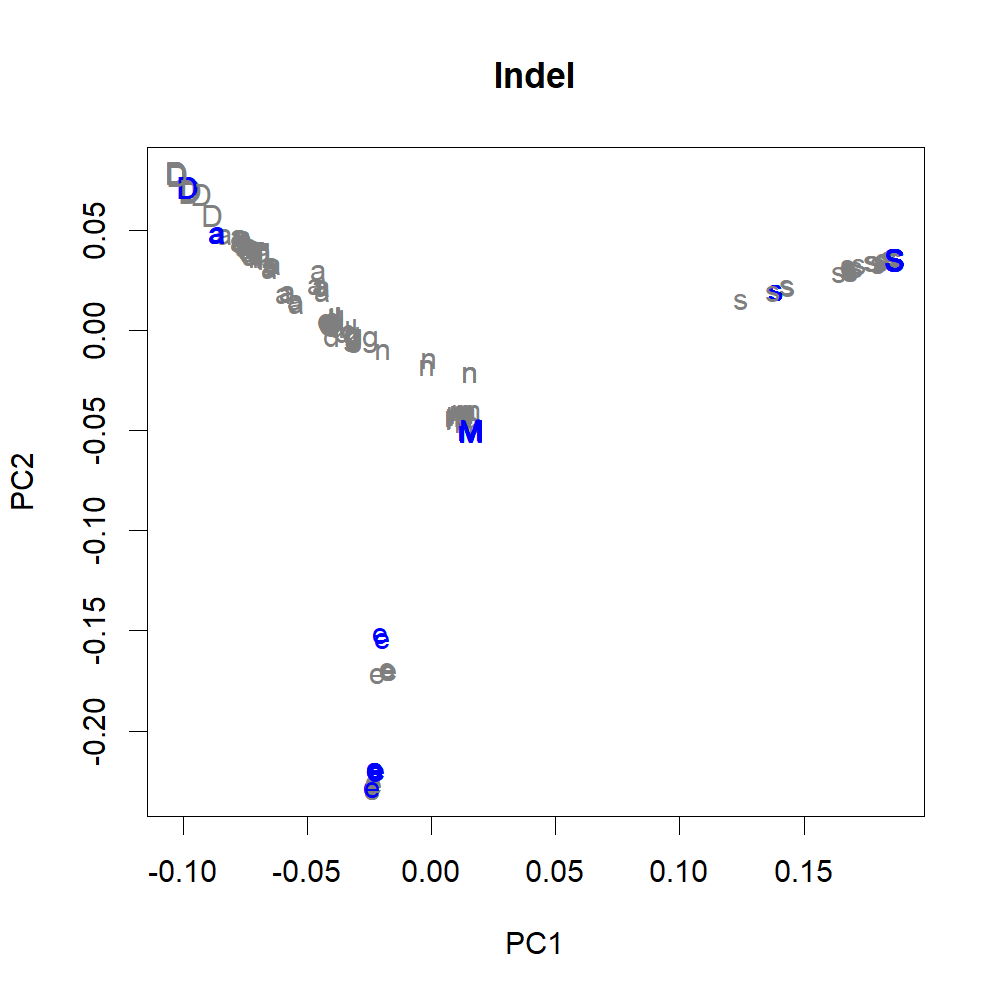

### S4b.png

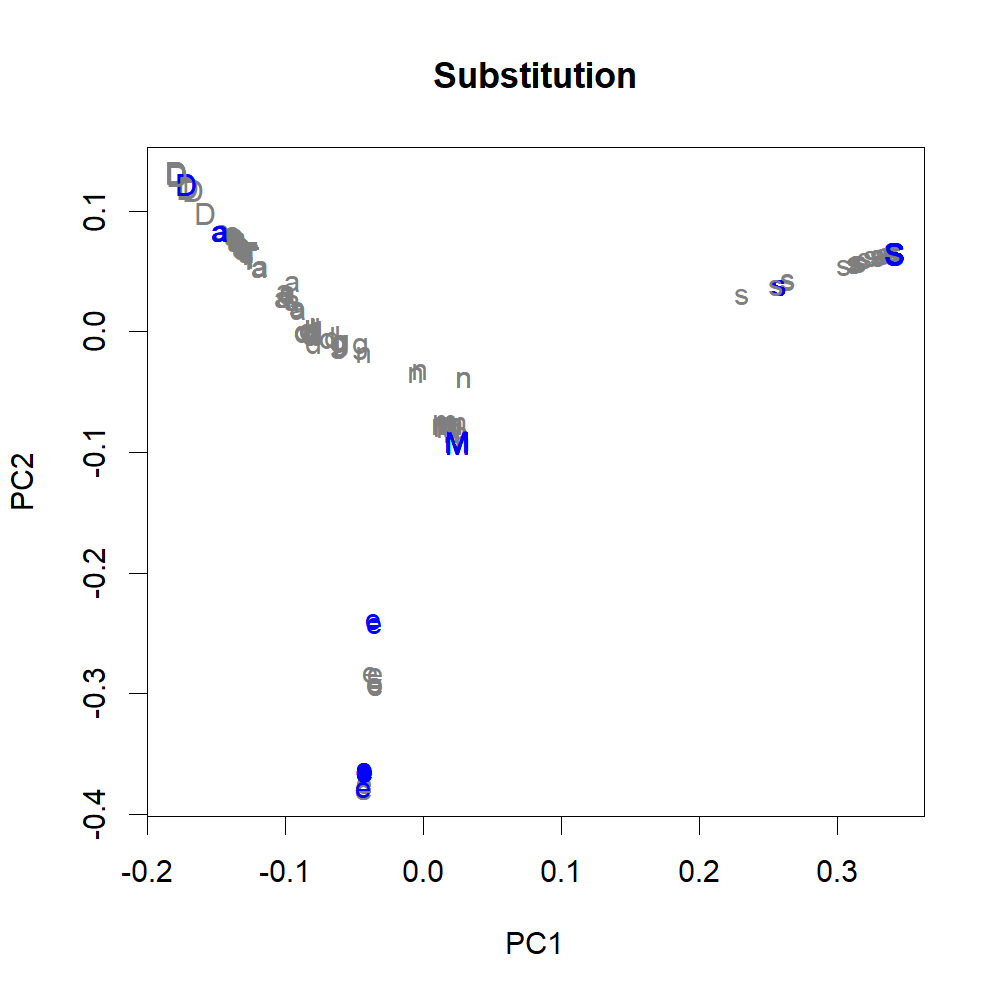

### S5a.png

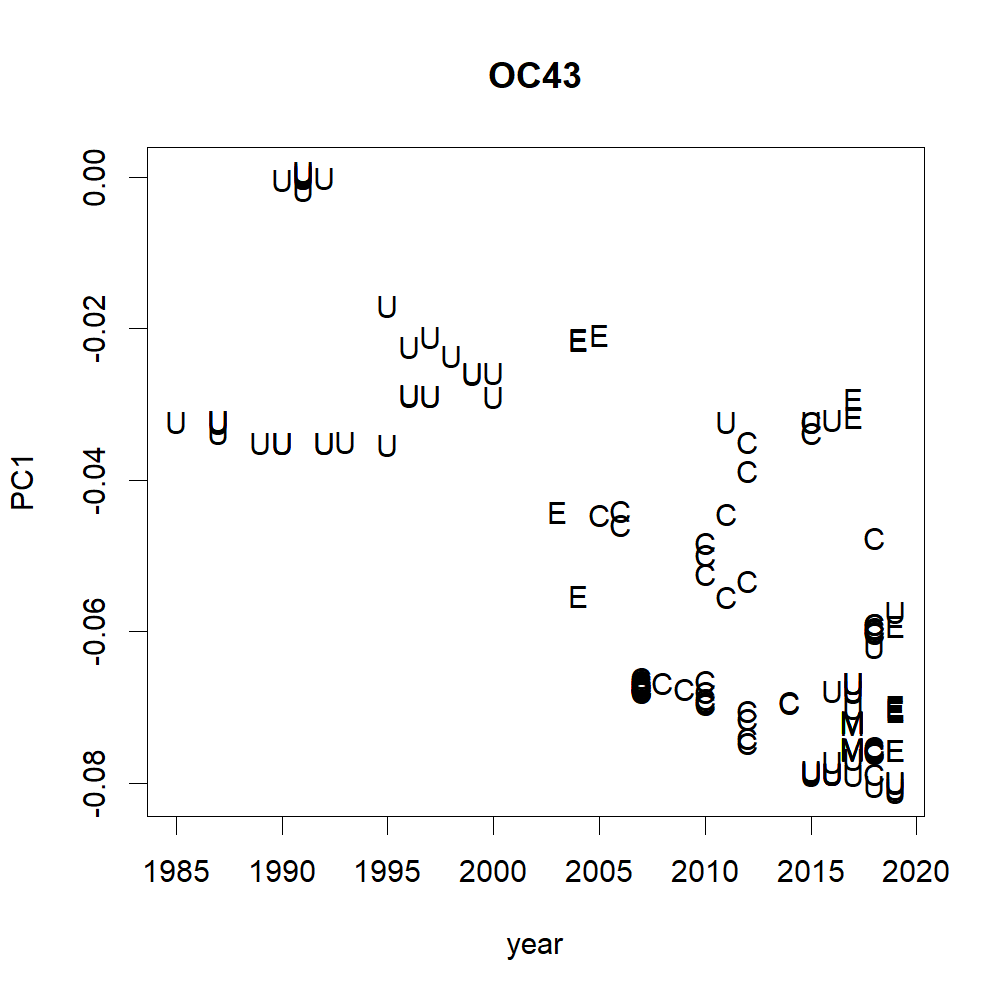

### S5b.png

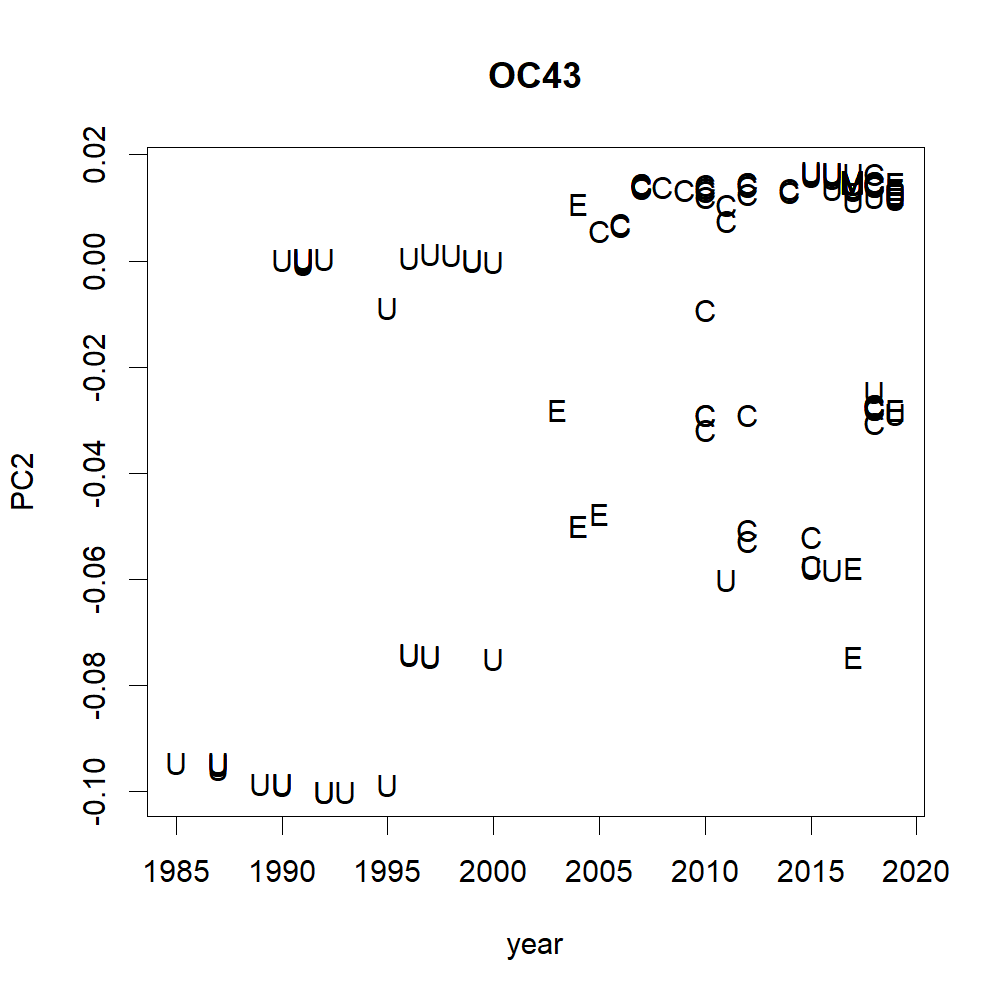

### S5c.png

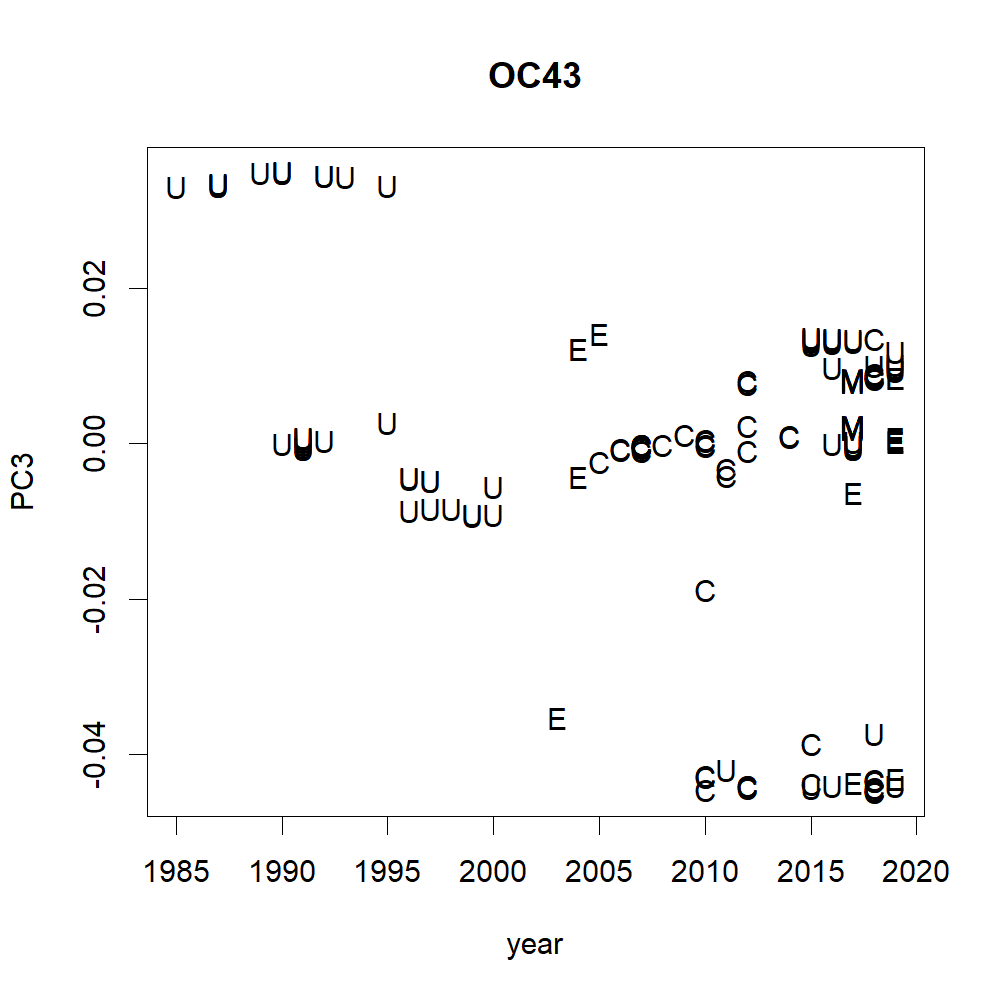

### S5d.png

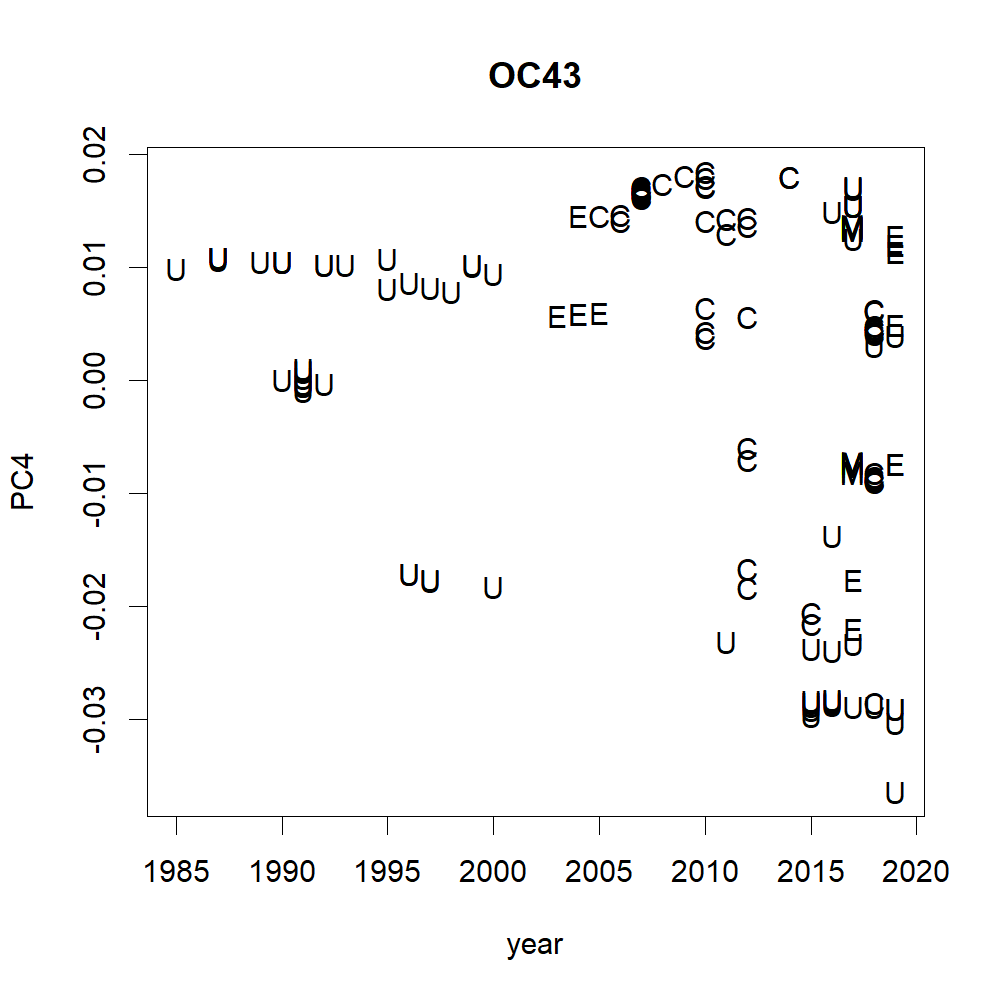

### S6a.png

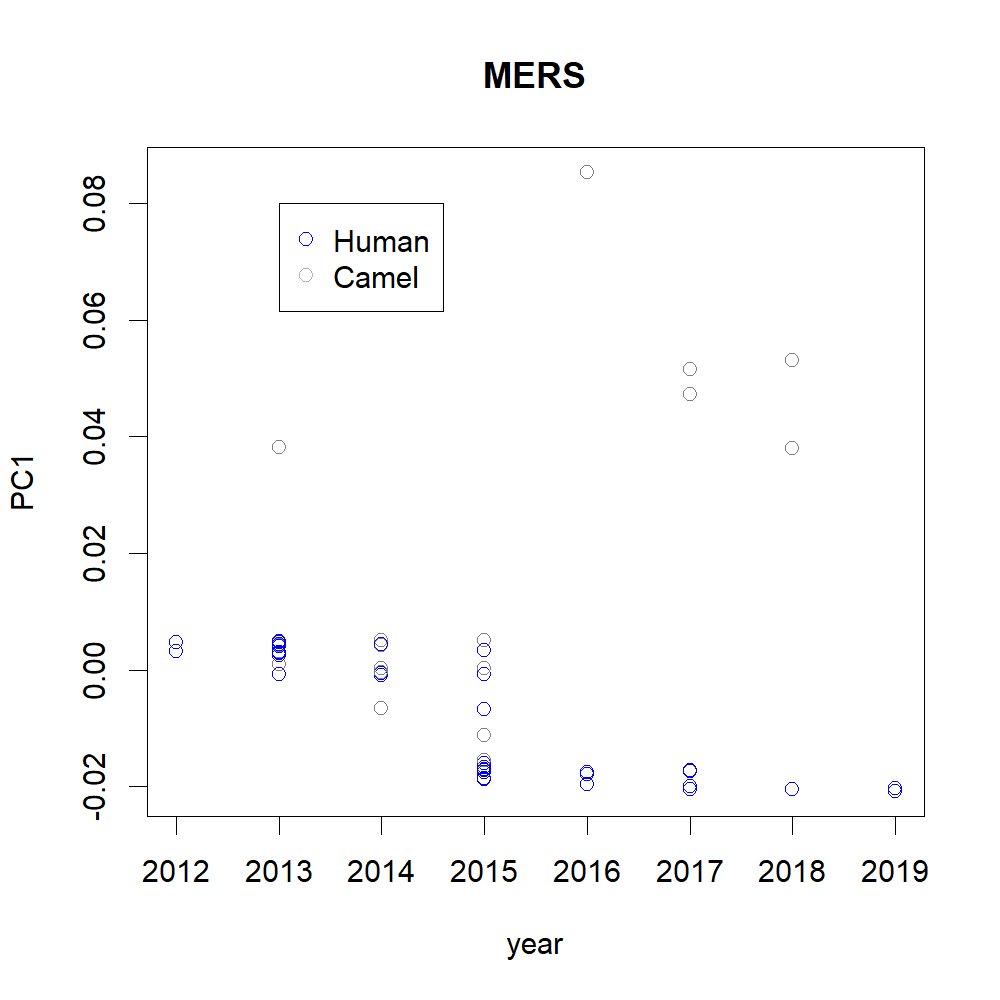

### S6b2.png

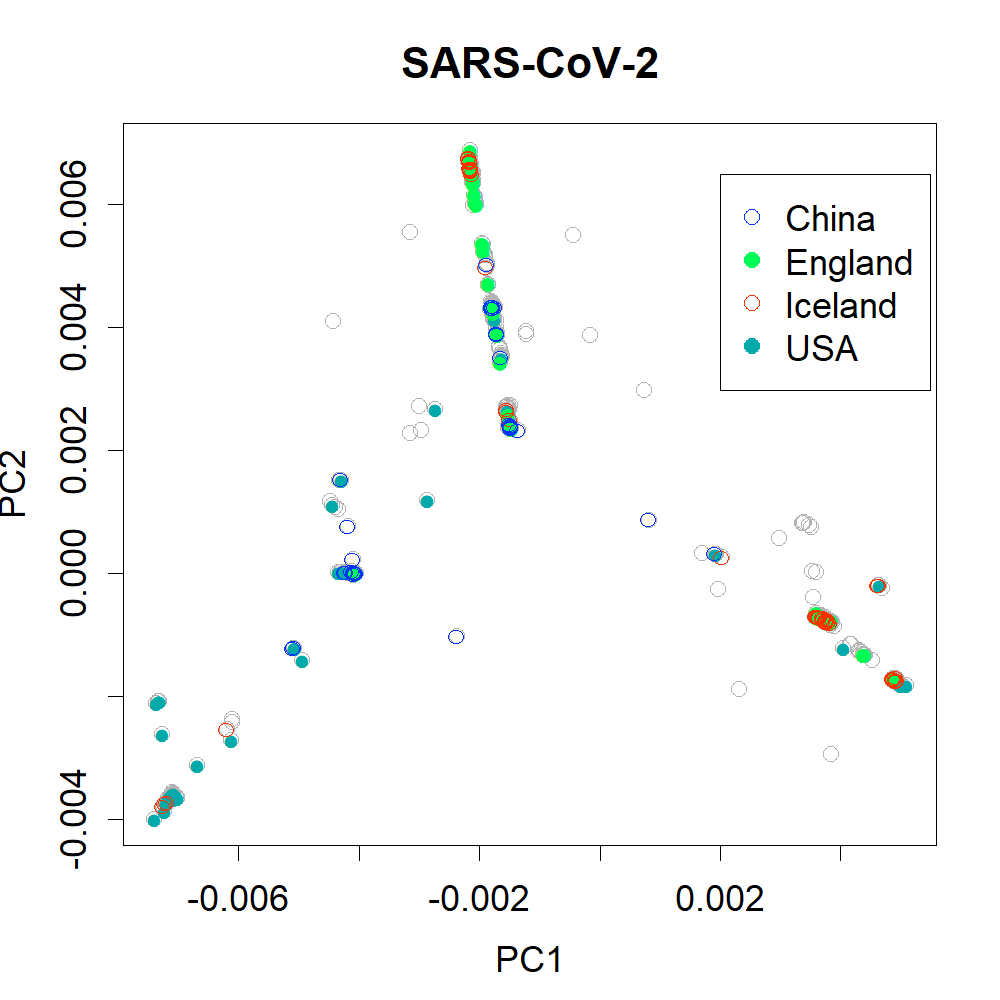

### S6b3.png

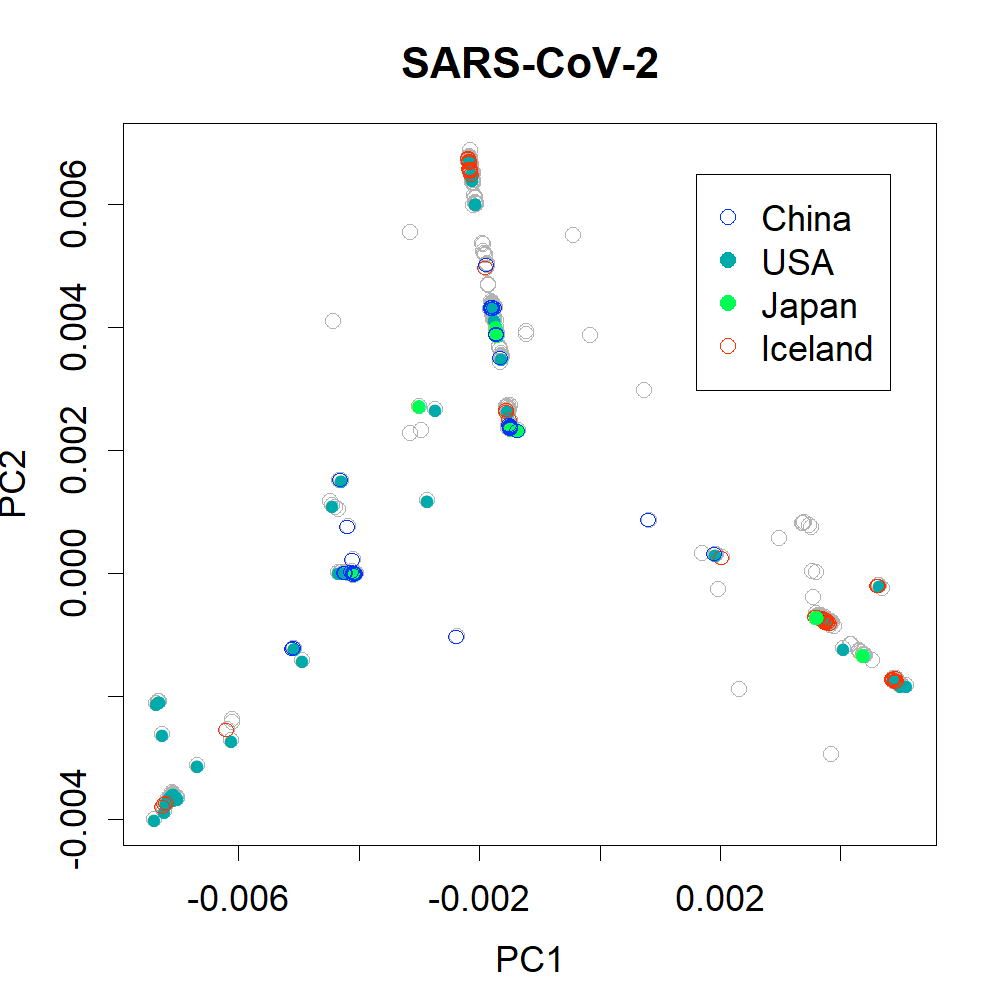

### S6b.png

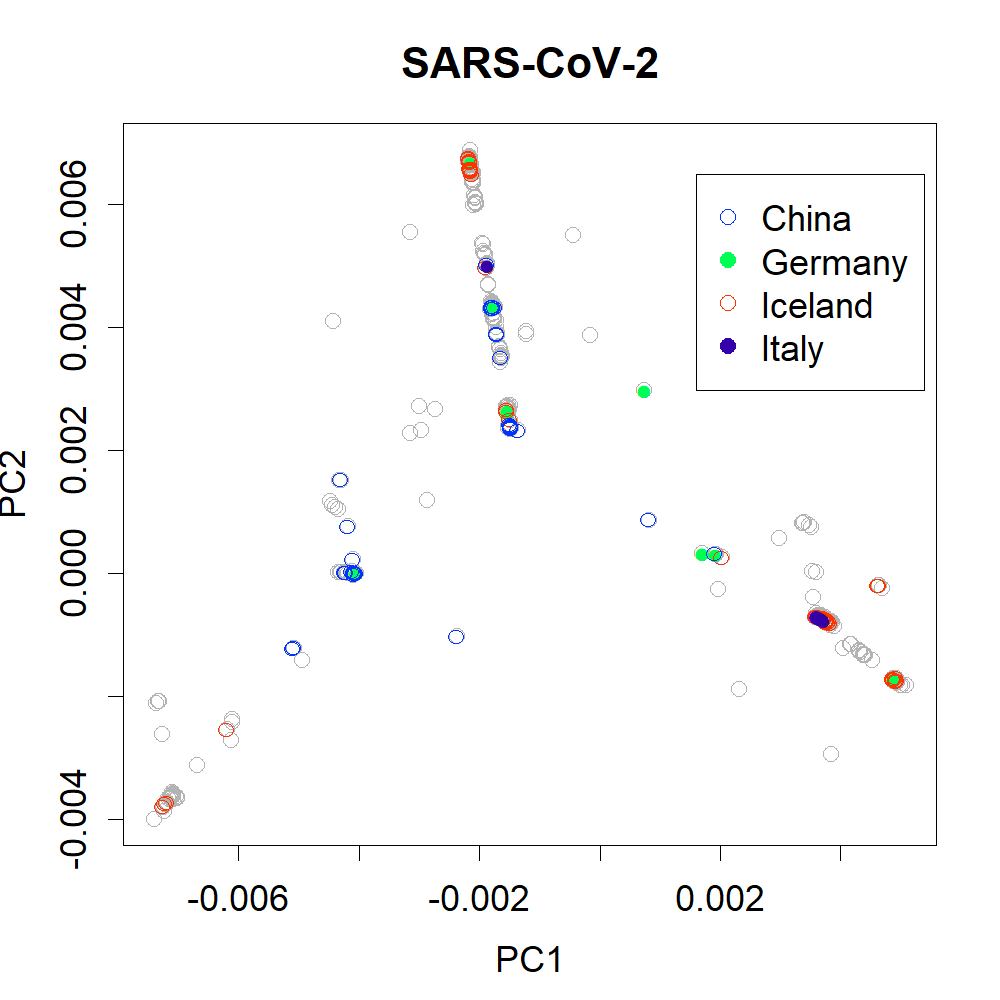
